# A benchmark of the somatic mutation landscape using single-cell and single-molecule whole-genome sequencing

**DOI:** 10.1101/2025.08.31.673349

**Authors:** Jacob E. Bridge, Chuwei Xia, Chen Zheng, Kristian Markon, Robert F. Krueger, David Masopust, Lei Zhang, Xiao Dong

## Abstract

Accurate detection of somatic mutations in noncancerous cells is critical for studying somatic mosaicism, a process implicated in aging and multiple chronic diseases. However, single-cell and single-molecule DNA sequencing platforms differ in their error profiles, coverage biases, and sensitivity to specific mutation types, complicating cross-platform comparisons. Here, we present *in vitro* and *in silico* benchmarks to quantify true-positive and false-positive rates in single-cell whole-genome sequencing using Single-Cell Multiple Displacement Amplification, and in single-molecule sequencing using Nanorate Sequencing (NS) and whole-genome NS (WGNS). Using standard cell lines, we show that all three methods detect single-nucleotide variants (sSNVs) and small insertions and deletions (sINDELs) with high accuracy, but differ in genomic coverage and susceptibility to artifacts. Method-specific biases influence mutational signatures and hotspot detection. Applying results of the benchmark to IMR-90 fibroblasts, we estimate higher *in vitro* mutation rates using NS than expected from *in vivo* data, consistent with potential replication stress and culture-associated DNA damage. Overall, our study highlights the substantial impact of sequencing platform-specific biases on somatic mutation detection and interpretation, and lays the foundation for standardized, cross-platform-comparable analyses of somatic mosaicism in normal human tissues.

## Introduction

Due to errors in DNA damage repair, somatic mutations inevitably accumulate over time. Somatic mutations in key functional genes drive ∼90% of human cancers, produce myriad genetic disorders, and have been linked to geriatric neurodegeneration, reduced muscle strength, and immune dysfunction^1–6^. Accurate quantification of somatic mutation burdens is essential for understanding the pathology of age-related diseases. Because DNA sequencing errors exceed the somatic mutation frequency, often 10^−7^ or less per base pair at the single cell level, by several orders of magnitude, conventional bulk DNA sequencing requires multiple variant-containing reads to distinguish mutations from technical artifacts. However, unlike in tumors or developing embryos, somatic variants in healthy adult tissues are frequently limited to the single-cell level. As a result, it is often impossible to obtain multiple variant-containing reads of mutations unique to individual cells using conventional bulk sequencing, even at sequencing depths in the 100x or 1000x range. To address this issue, two major categories of methods are under development: (i) single-cell DNA sequencing, which relies on accurate DNA amplification performed either *in situ* or *in vitro* to produce many variant-containing reads, e.g., natural clones^7^, SCMDA^8^, LIANTI^9^ and PTA^10^; and (ii) single-molecule (or “duplex”) sequencing of bulk DNA, which independently sequences both strands of the same DNA molecule, e.g., duplex-seq^11^, NanoSeq^12^, CODEC^13^, SMM-seq^14^, and HiDEF-seq^15^. The latter leads to more accurate mutation calling because, in principle, the probability that complementary technical artifacts occur at the same position on both strands is one third the square of the individual strand error rate. Both single-cell and single-molecule sequencing methods have contributed substantial innovations, including recent findings that thousands of mutations accumulate with age in every human cell, regardless of health status or tissue type^16^.

Continuous technical advancements remain essential for improving somatic mutation characterization. However, while new sequencing protocols can be validated against existing methods, such as single cell clones, there is no universal benchmark to assess the accuracy of somatic mutation calling at the single-cell level. The cell lines used for validation by different research groups differ widely by tissue type, donor age, and time in culture, resulting in distinct mutation burdens and signatures that make it difficult to gauge the relative performance of each method.

Validating somatic mutation calling at the single-cell level is further complicated by the lack of “ground truth” variants. Because each somatic mutation is unique to a parent cell and its progeny, the true frequency and distribution of mutations within any given cell is unknown. Without this information, while it is possible to infer the fidelity of a mutation characterization pipeline through agreement with existing methods, the performance metrics of each method cannot be reliably determined. As a result of these factors, conclusions about somatic genome mosaicism and its impact could be substantially confounded by choice of pipeline, at the expense of biological significance.

To more thoroughly evaluate current and future mutation characterization pipelines, we generated somatic mutation profiles for two model B lymphoblastoid cell lines using state-of-the-art single-cell whole-genome and single-molecule sequencing methods^8,12,17,18^. We also developed *in silico* and *in vitro* approaches to establish a ground truth mutation burden, using the former to identify the mutation rate per cell division in human lung fibroblasts. The methodologies and findings outlined in this study will serve as a robust technical foundation for future analyses of somatic mutations in non-cancerous tissues.

## Results

### Deep single-cell whole-genome and single-molecule sequencing of two human lymphoblastoid benchmark lines

In 2016, the National Institute of Standards and Technology (NIST) established several cell lines as benchmark reference materials^19^, which have become widely used to validate new DNA sequencing applications^20–22^. To facilitate comparisons of library preparation and mutation calling methods, we established two of these cell lines– GM12878 (G1) and GM24385 (G2)–as benchmarks for somatic mutation calling.

We obtained G1 and G2 cells, along with their genomic DNA, from the Coriell Institute. For single-cell whole-genome sequencing, we amplified individual lymphoblastoid genomes using Single-Cell Multiple Displacement Amplification^8,17^ (SCMDA; n = 5 cells/line), performed whole-genome sequencing using Illumina 150bp paired-end reads (mean depth per cell = 42.1 ± 7.4x; **Table S1**), and called somatic single-nucleotide variants (sSNVs) and somatic small insertions and deletions (sINDELs) using sccaller (v2.0). For single-molecule sequencing, we applied Nanorate Sequencing^12^ (NS; n = 3 replicates/line; mean WGS-equivalent depth = 56.5 ± 5.5x), as well as a newly updated protocol for whole-genome NS^18^ (WGNS; n = 1 replicate/line; mean depth = 36.8 ± 2.6x), to bulk genomic DNA. For standard NS, we replaced the original ligation mix with a NEB Ultra II Ligation Module to increase library yield (Materials and Methods). We used conventional bulk Whole-Genome Sequencing (WGS; mean depth = 42.6 ± 3.9x) data from each line (n = 1) as a matched normal to screen out germline variants, allowing us to call somatic mutations from SCMDA, NS and WGNS data (**Fig. 1**).

**Figure 1.**
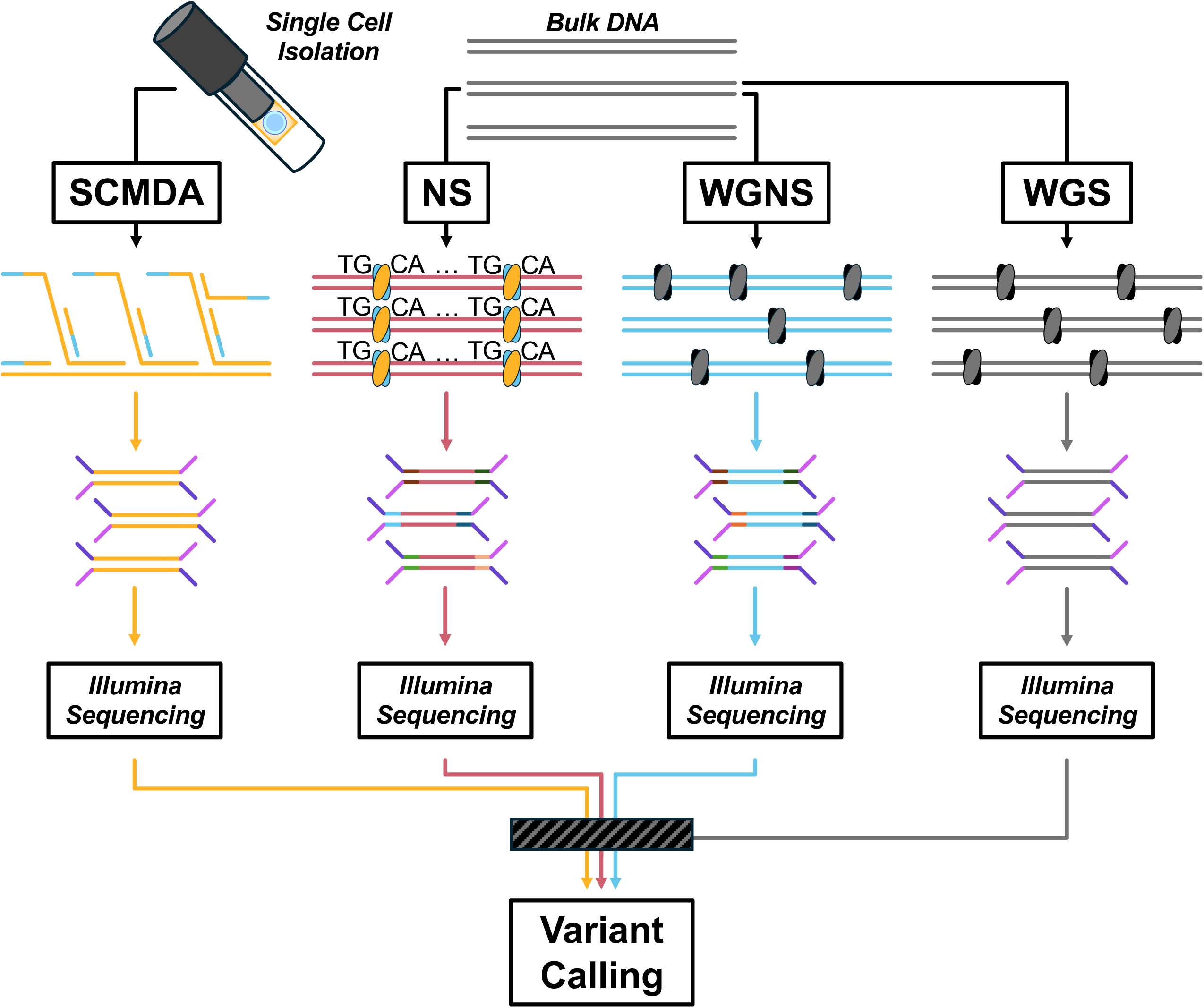
Overview of single-cell and single-molecule sequencing pipelines used to detect somatic mutations. Single cells were subjected to SCMDA. Bulk DNA from the same cell lines was subjected to standard WGS, NS, or WGNS. All libraries were subjected to Illumina sequencing, then analyzed based on their sequencing characteristics. We also performed variant calling on the SCMDA, NS, and WGNS libraries, using WGS data as a matched normal to screen out germline variants.

### Genomic coverage and strand bias

Before assessing mutation calling performance, we investigated the sequencing characteristics of each pipeline. We began by comparing the sequencing coverage of NS, WGNS and SCMDA to that of WGS. After downsampling each pipeline’s raw sequencing reads to a mean depth of 30x, we found that NS achieved a maximum potential coverage of ∼60% (**Fig. 2A**), reflecting the limitation of restriction enzyme digestion to conserved 5’-TGCA-3’ sites. As expected, WGNS, SCMDA, and WGS exhibited a maximum potential coverage of ∼90%. Based on the pipelines’ Lorenz curves (**Fig. 2B**), we found that WGNS exhibited a more even read distribution than NS or SCMDA. We also examined the fraction of bases covered at different sequencing depths for each sample (**Figs. 2C & S1A**). In agreement with the Lorenz curve data, NS exhibited the most biased read distribution, followed by SCMDA, WGNS, and WGS. Unlike the other sequencing methods, NS also displayed an exponential decline in coverage with increasing depth and lacked a local maximum in the 15-35 read range, once again aligning with restriction enzyme digestion at conserved target sites.

**Figure 2.**
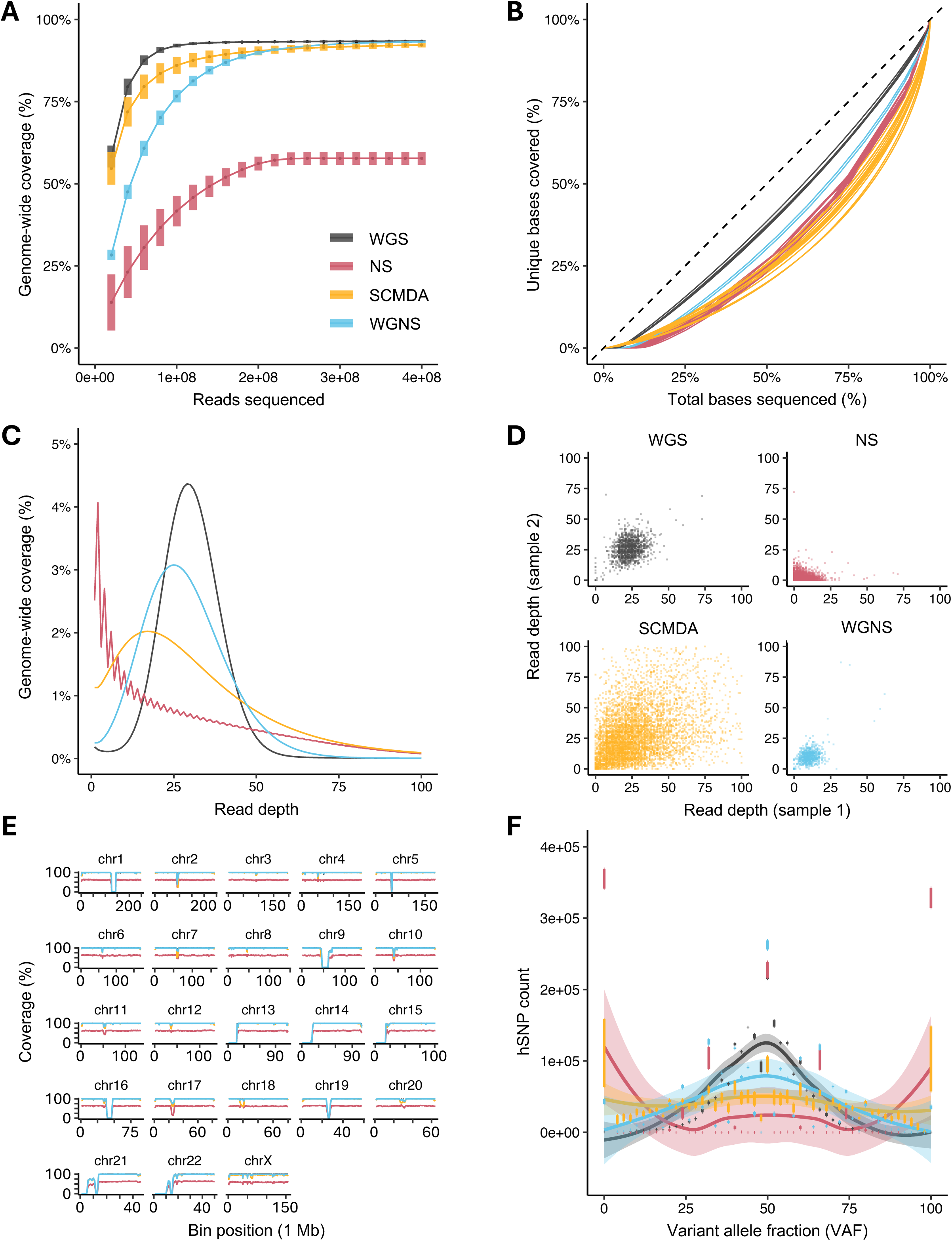
Sequencing characteristics of single-cell and single-molecule pipelines. (A) Sequencing coverage as a function of read count in WGS, NS, SCMDA, and WGNS. For each method, reads within each replicate were resampled, and coverage was calculated as fraction of the reference genome. Data for GM12878 (G1), GM24385 (G2), and IMR-90 samples are pooled, and displayed as the mean ± standard deviation across replicates. n = 6 WGS, 10 NS, 10 SCMDA, and 2 WGNS samples. (B) Lorenz curve of sequencing uniformity. For each replicate, the cumulative fraction of sequenced bases was plotted against the fraction of unique covered bases, relative to the total coverage of the replicate. The dashed line represents uniform coverage across all sequenced bases. (C) Coverage-depth distribution for each pipeline. Each replicate was downsampled to a mean sequencing depth of 30x following PCR deduplication. For each method, lines represent the average fraction of genome covered at a given read depth across replicates. The jagged line for NS arises from the deduplication algorithm, which assigns a depth of two to any read family containing at least one top and bottom strand duplicate, thereby favoring even read depths. Coverage fractions at a read depth of zero are not displayed, but are included in Fig. S1. (D) Inter-sample bias in read depth. Within each method, pairwise combinations of replicates were selected, then evaluated for read depth at 100 random positions. Due to the low number of replicates, 1000 random positions were used for WGNS. Points represent individual sample pairs, with read depths limited to a maximum of 100x. (E) Fraction of bases covered within 1 Mb bins across autosomes and chromosome X. For each method, lines represent the mean coverage across all replicates. (F) Allelic bias in germline heterozygous SNPs (hSNPs). For each replicate, the total number of hSNP sites exhibiting a given variant allele fraction was plotted, using a VAF bin size of 2%. Individual bars display the mean ± standard deviation across replicates, while the dark lines and shaded regions represent a smooth fit for each method. Only samples from G1 and G2 (n = 1 WGS, 3 NS, 5 SCMDA, and 1 WGNS samples per cell line) are included in Fig. 2F.

Next, we tested whether sequencing depth at randomly selected loci remained consistent across samples prepared with the same pipeline. While locus-specific read depth was largely homogenous within pairs of WGS and WGNS samples (**Fig. 2D**), SCMDA was less uniform, suggesting stochastic genome amplification bias between different single cells, while NS displayed the least consistent coverage between samples. NS also produced fewer unique duplex-covered bases than WGNS, sequencing less than 30% of the genome at sufficient depth for mutation calling, while sporadically covering another 35-40% of the genome (**Table S2**). This equated to unique duplex coverage efficiencies, relative to an unbiased Poisson distribution, of only 30-40% for NS, compared to >90% for WGNS. After examining the chromosome-wide coverage of each protocol, we found that WGS, SCMDA, and WGNS covered all genomic loci except centromeres (**Fig. 2E**). Although NS exhibited similar chromosome-wide coverage patterns, the maximum coverage within any 1 Mb-bin did not exceed 70%.

Finally, we assessed the allelic bias of each protocol using germline heterozygous single-nucleotide polymorphisms (hSNPs; **Fig. 2F**). As expected, all four protocols produced hSNP variant allele fractions (VAFs) centered around 50%, while WGNS, NS, and SCMDA exhibited greater bias than WGS.

Overall, these results demonstrate that while WGNS and SCMDA provide robust whole-genome coverage, NS is limited to smaller fragments of the genome. As a result, NS may not be an ideal approach to investigate mutation accumulation in particular functional regions, such as the exons of cancer driver genes.

### Accuracy of somatic mutation calling varies by method

Historically, the absence of ground truth somatic mutation data has made it difficult to distinguish true positives from technical artifacts, limiting assessments of mutation calling performance. To address this issue, we developed an *in silico* approach to define “pseudo-somatic” mutations within sequencing data. Rather than using WGS data from the same subject as a matched normal for somatic mutation calling, we supply WGS data from a different genetic background, such as another human subject, or, in the case of a mouse study, a distinct strain (**Fig. 3A**). In this case, we screened SCMDA, WGNS, and NS datasets from G1 samples against WGS data generated from G2 bulk DNA, and *vice versa*. We reasoned that resulting somatic mutation calls could arise from one of three sources: (i) real somatic variants within the G1 NS, WGNS, or SCMDA samples; (ii) technical artifacts; or (iii) germline variants within G1 NS, WGNS, or SCMDA samples that are not present in G2 WGS samples. We limited our analysis to germline hSNPs and hINDELs, reflecting the exceedingly low likelihood of possessing homozygous somatic mutations. In humans, germline variants are three orders of magnitude more common than somatic variants and technical artifacts combined (**Figs. 3B-C**). As a result, we assumed that mutation calls matching source (iii) were “pseudo-true positives” (pTPs), and denoted all other mutation calls as “pseudo-false positives” (pFPs). This approach allowed us to approximate the performance metrics of somatic mutation calling.

**Figure 3.**
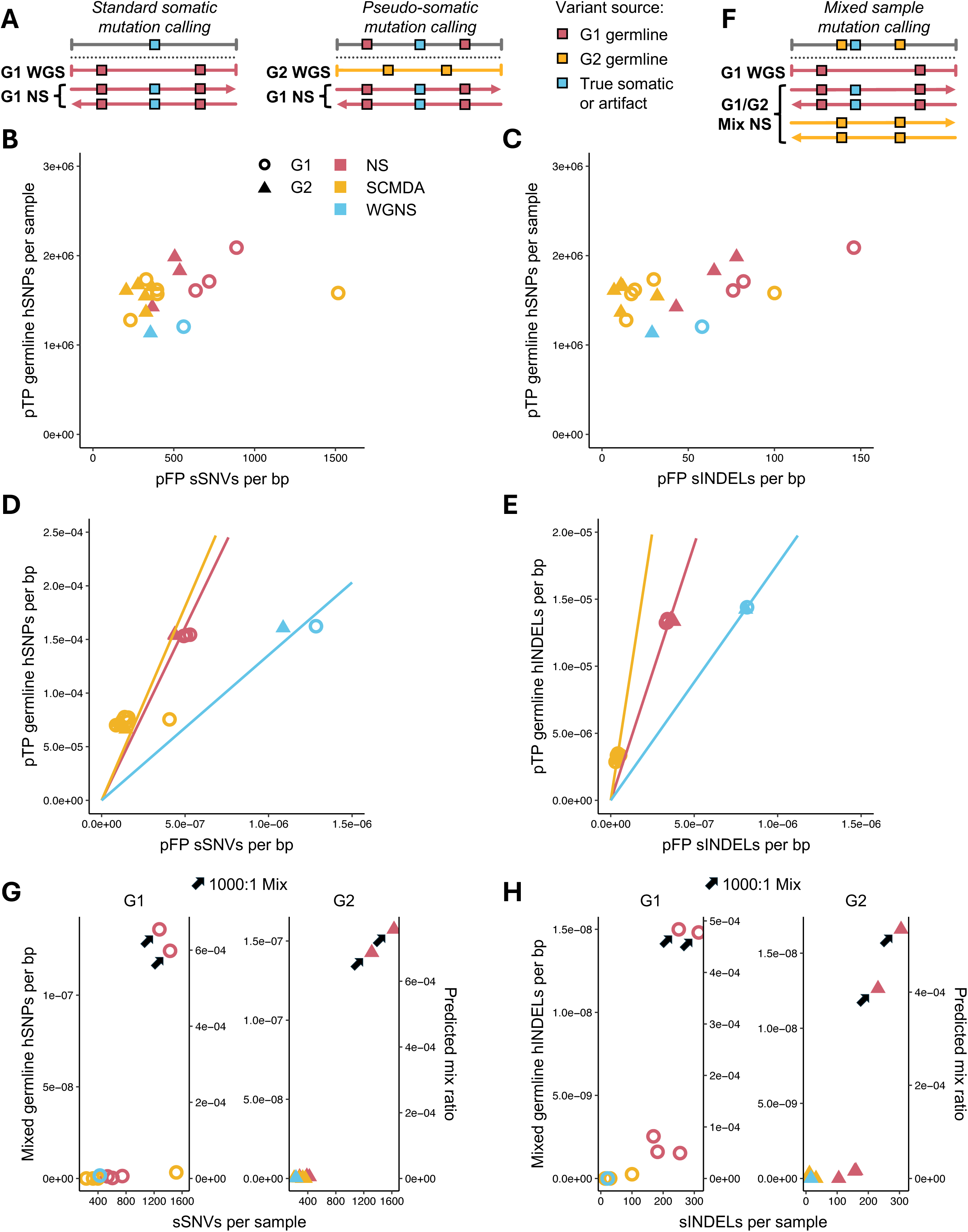
*In silico* and *in vitro* assessment of somatic mutation calling accuracy. (A) Visual representation of *in silico* pseudo-somatic mutation calling. Unlike standard mutation calling, pseudo-somatic mutation calling uses WGS data from a different genetic background as a matched normal, causing germline variants in NS, SCMDA, and WGNS libraries to appear as “pseudo-somatic” mutations. Pseudo-true positive (pTP) variants align with known germline mutations, while pseudo-false positive (pFP) variants include true somatic mutations and technical artifacts. (B-C) Total pFP somatic SNVs (sSNVs) and somatic INDELs (sINDELs) and pTP germline hSNPs and germline heterozygous INDELs (hINDELs) detected in G1 and G2 samples. Each point represents the mutation counts of a single technical replicate. n = 3 NS, 5 SCMDA, and 1 WGNS samples per cell line. (D-E) Per base pair (bp) mutation burdens of samples displayed in Figs. 3B-C. Slopes of the solid lines indicate the average ratio of pTP to pFP variants detected by each pipeline. A larger slope indicates higher accuracy. (F) Visual representation of *in vitro* mixed germline mutation calling. NS libraries are prepared and sequenced from 1:1000 mixtures of bulk DNA from two different cell lines, then called against WGS data from the 1000x cell line. Germline variants in the 1x cell line, as well as true somatic variants and technical artifacts from both cell lines, will be called as somatic mutations. (G-H) Mixed germline hSNPs and hINDELs from 1x cell lines as a function of total detected variants. Predicted mix ratios were calculated by dividing observed germline mutations per bp by the known frequency of germline mutations within the 1x cell line. The “G1” and “G2” labels at the top of each graph indicate the 1000x cell line, with the other serving as the 1x cell line. Mixed samples are indicated by arrows. Data from unmixed samples (Figs. 3B-E) is included for comparison. n = 2 replicates per mixture.

Before assessing pipeline accuracy, we first determined the total number of sSNVs and sINDELs called by SCMDA, NS, and WGNS. Because the common SNP and noise masks used for NS and WGNS could be applied to any sequencing pipeline, we counted both “masked” and “pass” calls as somatic mutations during pipeline comparison (see Materials and Methods). At mean sequencing volumes of 4.9 ×10^8^, 3.9 ×10^8^, and 3.8 ×10^8^ read pairs (**Table S1**), respectively, SCMDA, NS, and WGNS detected an average of 437, 609, and 459 sSNVs per sample (**Fig. 3B**), as well as 25, 82, and 44 sINDELs per sample (**Fig. 3C**).

We then replaced each pipeline’s matched normal with WGS data from the opposite cell line, and quantified the number of pTPs and pFPs per base pair (bp) called by SCMDA, NS, and WGNS. For both hSNPs and hINDELs (**Fig. 3D-E**, in which a larger slope indicates better accuracy as demonstrated also in **Fig. S2**), NS and WGNS detected far more pTPs than SCMDA. Based on the expected germline mutation burdens of each cell line, the average sensitivities of SCMDA, NS, and WGNS, respectively, were 35.1%, 74.4%, and 78.0% for hSNPs and 10.5%, 43.9%, and 47.0% for hINDELs. However, SCMDA displayed far greater precision than the other pipelines, reporting 3.1-fold fewer pFP sSNVs/bp and 8.9-fold fewer pFP sINDELs/bp than NS, as well as 7.8-fold fewer pFP sSNVs/bp and 20.8-fold fewer pFP sINDELs/bp than WGNS. Based on each pipeline’s ratio of pTPs to pFPs, NS and SCMDA were similarly effective at calling SNVs, while SCMDA was by far the most reliable for INDEL calling. Despite exhibiting the highest sensitivity for both hSNPs and hINDELs, WGNS was consistently the least reliable pipeline for variant calling, due to its disproportionately high error rate.

To ensure mismatched germline variants are detectable *in vitro* as well as *in silico*, we prepared NS libraries from 1:1000 mixtures of G1 and G2 genomic DNA. After sequencing these libraries, we called sSNVs and sINDELs against WGS data from the 1000x cell line, causing heterozygous germline variants from the 1x cell line to appear as pseudo-somatic mutations (**Fig. 3F**). Because it requires bulk DNA inputs, this method is suitable for testing single-molecule sequencing protocols, but is not compatible with single-cell protocols such as SCMDA. Nevertheless, we found that NS could clearly distinguish mixed samples from pure samples based on 1x line hSNP and hINDEL burdens (**Figs. 3G-H**). Furthermore, based on an expected mix ratio of 1:1000, NS exhibited a mean hSNP sensitivity of 67.4% and a mean hINDEL sensitivity of 47.7%, closely matching the results of our *in silico* model.

Overall, our *in vitro* test confirmed that heterozygous germline variants are a robust source of ground truth “pseudo-somatic” mutations, reinforcing our *in silico* approach as a reliable tool to assess the performance metrics of pipelines used for somatic mutation characterization. SCMDA exhibited the greatest precision for both pseudo-sSNVs and pseudo-sINDELs, while NS and WGNS balanced increased sensitivity with higher pFP rates.

### Somatic mutation burdens of two human lymphoblastoid benchmark lines

In addition to developing our pipeline validation approaches, we quantified the sSNV and sINDEL burdens of the G1 and G2 cell lines using single-cell and single-molecule sequencing, establishing a benchmark for future methods development. For this analysis, we compared SCMDA, NS, and WGNS data from unmixed samples to WGS data from the corresponding cell line. Notably, for each computational pipeline, we followed the steps and parameter settings outlined in its original publication. As a result, the variant calling parameters used for NS and WGNS were not identical (see Materials and Methods), causing several artificial mutational hotspots to appear in the WGNS data (see below). To further reduce the false positive rate of NS and WGNS, we also implemented the SNP and noise masks described in their respective protocols. In exchange for masking 2.25% of the genome, NS reported 29.4% fewer sSNVs and 48.7% fewer sINDELs, while WGNS reported 37.9% fewer sSNVs and 53.4% fewer sINDELs (**Table S3**). Across all three pipelines, we detected a total of 7,549 sSNVs and 532 sINDELs. On a per-sample basis, individual pipelines produced the following average mutation counts, for G1 and G2 respectively: SCMDA reported 574.4 and 300 sSNVs, as well as 36 and 14.6 sINDELs; NS reported 550.3 and 317 sSNVs, as well as 46.3 and 34 sINDELs; and WGNS reported 363 and 212 sSNVs, as well as 22 and 16 sINDELs (**Table S3**).

We then calculated the sSNV burden for each sample as the ratio of sSNVs to total sequenced bases, adjusting for estimated sSNV calling accuracy using *in silico* pTPs and pFPs (Materials and Methods; **Fig. S3A**; **Table S3**). G1 (mean = 4.3, 1.9, and 1.9 ×10^−7^/bp for SCMDA, NS, and WGNS, respectively) exhibited a higher sSNV burden than G2 (mean = 2.4, 1.1, and 1.2 ×10^−7^/bp, respectively) across all three pipelines (**Fig. 4A**). Although SCMDA reported ∼2-fold higher mean sSNV burdens than the other pipelines, none of these differences were statistically significant (ANOVA with Tukey’s HSD, 0.055 ≤ P ≤ 0.743). We also observed greater variation in SCMDA replicates compared to NS samples, likely because single-cell sequencing captures variation among individual cells, while NS produces an “assembly” of mutation profiles collected from fragments of many genomes. For sINDELs (**Fig. 4B, Fig. S3B**), G1 (mean = 9.0, 2.8, and 1.9 ×10^−8^/bp for SCMDA, NS, and WGNS, respectively) again exhibited a higher mutation burden than G2 (mean = 3.7, 1.9, and 1.4 ×10^−8^/bp, respectively). In line with sSNV data, SCMDA detected higher sINDEL burdens than the other pipelines, although again these differences were not significant (0.457 ≤ P ≤ 0.620). Inconsistencies in mutation burden between single-cell and single-molecule sequencing methods may have resulted from SCMDA’s small sample size, differential application of SNP and noise masks, and remaining bias in pTP and pFP estimation (see discussion).

**Figure 4.**
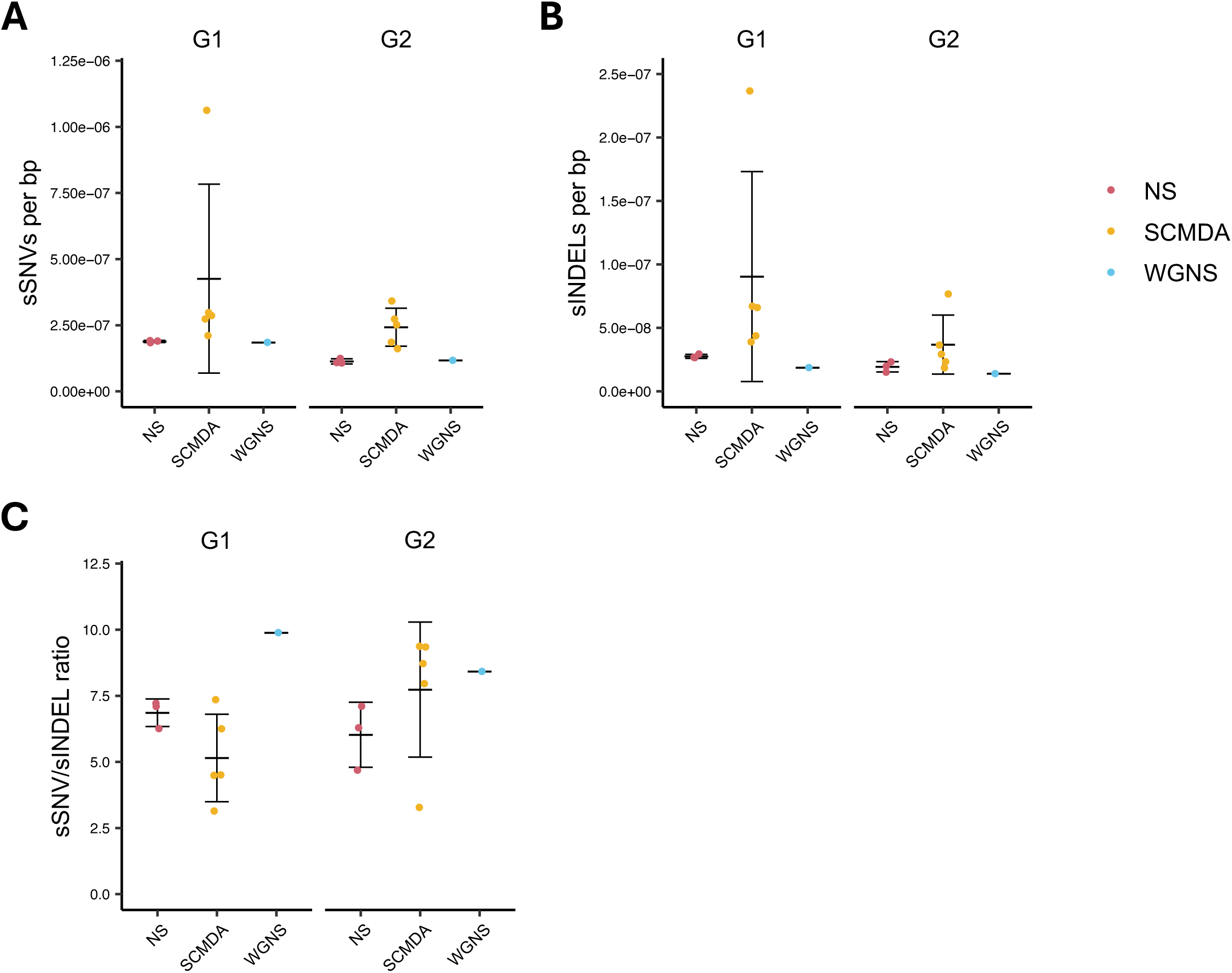
*In silico*-adjusted mutation burdens in lymphoblastoid cell lines. (A-B) sSNVs and sINDELs per bp detected in G1 and G2 samples by NS, SCMDA, and WGNS. Raw mutation burdens were multiplied by a correction factor derived from the pFP and pTP rates of each pipeline (see Materials and Methods). Each point represents a distinct technical replicate, while error bars indicate the mean ± standard deviation for each group of samples. n = 3 NS, 5 SCMDA, and 1 WGNS samples per cell line. (C) Ratios of sSNVs to sINDELs detected by each pipeline, calculated from the sSNV and sINDEL burdens displayed in Figs. 4A-B.

Within all three methods, the relative ratio of sSNVs to sINDELs remained largely consistent between G1 and G2 (**Fig. 4C**). All three pipelines yielded mean sSNV-to-sINDEL ratios between 5:1 and 10:1, which were similar to the germline mutation ratios of G1 and G2 (6.6:1 and 7.0:1 for cell line-specific heterozygous variants; **Table S4**). Unlike in the case of mutation burden, sSNV-to-sINDEL ratios produced by SCMDA did not differ noticeably from those of NS or WGNS. Overall, we found that following *in silico* adjustment, all three pipelines produced similar estimates of sSNVs/bp, sINDELs/bp, and sSNV-to-sINDEL ratio in G1 and G2 cell lines, providing a benchmark for testing novel methods of somatic mutation detection.

### In vitro mutation rate per cell division in human fibroblasts

Benchmarking and error correction allow for more robust assessments of somatic mutation burden, leading to improved characterization of physiological processes. One subject of particular interest is the somatic mutation rate per cell division. Given the established role of somatic mutations in tumorigenesis, as well as increasingly strong links between somatic mutations and aging, it is critical to determine the rate of mutation accumulation in human cells. Previously, we calculated the somatic mutation rate in adult dermal fibroblasts as a function of estimated post-zygotic cell divisions^23^. However, since the exact number of stem cell divisions prior to differentiation was unknown, these estimates may have included additional errors beyond those introduced during mutation calling.

We addressed this issue by performing *in silico*-adjusted NS on single-cell clones, using clone WGS data as a matched normal (**Fig. 5A**, *middle right*). Under this design, only *de novo* mutations that arose during clonal expansion would produce somatic mutation calls. Rather than using immortalized cell lines such as G1 and G2, we derived our clones from early-passage IMR-90 cells, a widely used primary human fibroblast line. Three IMR-90 single-cell clones were expanded to 5.7-7.8×10^6^ cells over 30 days, subjected to NS, then called for somatic mutations against clone WGS data. All three clones exhibited similar adjusted sSNV and sINDEL burdens, averaging 9.92 ×10^−8^ and 1.99 ×10^−8^/bp, respectively (**Figs. 5B-C**). This 5:1 sSNV-to-sINDEL ratio was similar to that observed in lymphoblastoid cell lines, but was far lower than the ratio detected in IMR-90 clone WGS libraries (**Figs. S4A-B**), suggesting that human cells experience disproportionately high sINDEL accumulation during *in vitro* culture. Stricter variant calling criteria for sINDELs relative to sSNVs may have also contributed to the Iower sSNV-to-sINDEL ratios observed in clone data.

**Figure 5.**
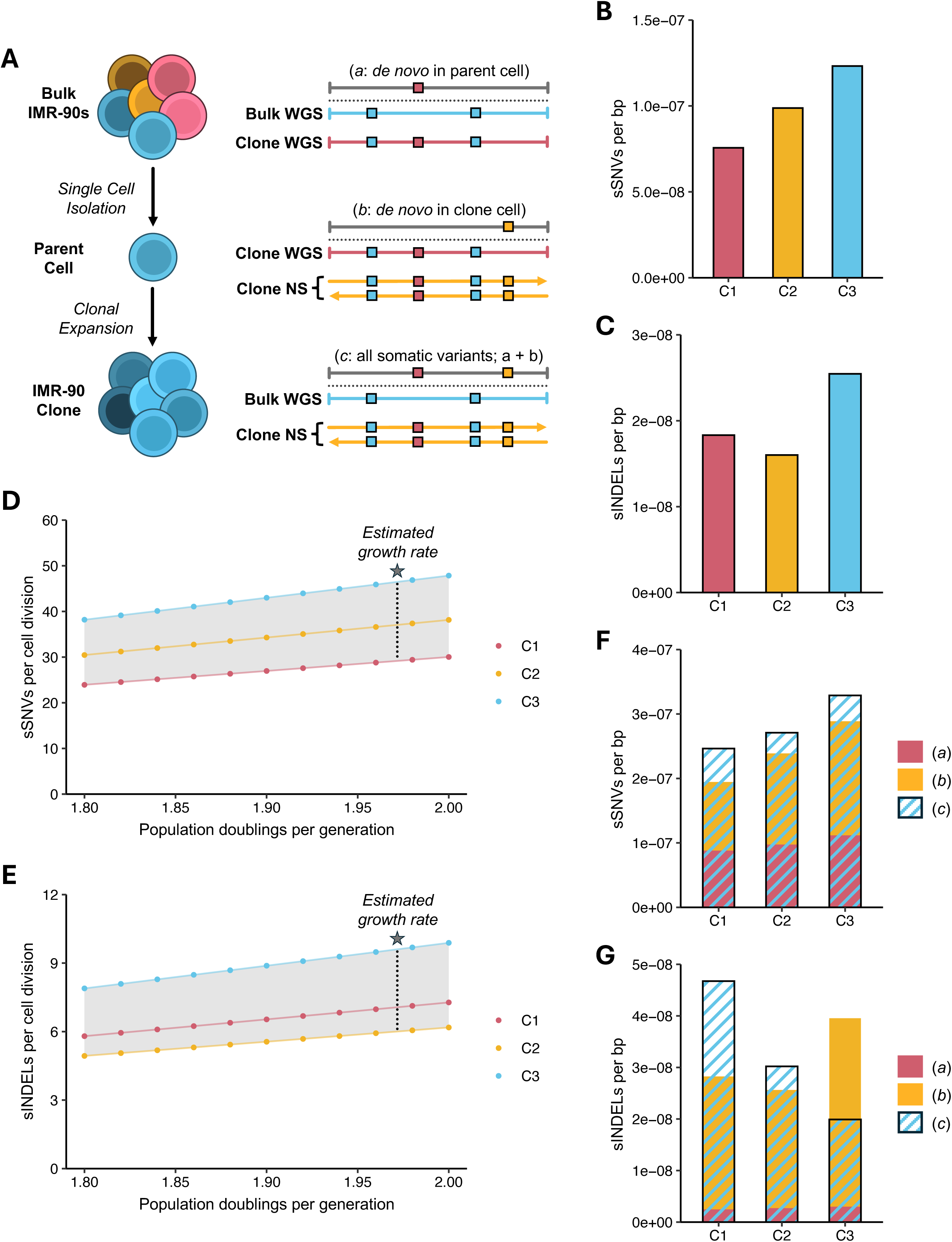
Quantifying mutation rate per cell division using *in silico*-adjusted NS. (A) Application of NS to clonal populations. Individual IMR-90 fibroblasts were isolated, then expanded into large clones over 30 days. IMR-90 clones, as well as a separate population of bulk IMR-90 cells, were then subjected to WGS and NS. Distinct combinations of bulk WGS, clone WGS, and clone NS data provide different selections of variants, which can be used to estimate mutation rate per cell division and validate NS. (B-C) sSNVs and sINDELs per bp detected in IMR-90 clones. Mutation burdens were calculated by screening clone NS libraries against clone WGS data (Fig. 4A, *middle right*), isolating variants that arose *de novo* during clonal expansion. A single NS and WGS library were constructed for each clone. n = 3 clones. (D-E) Mutation rate per cell division in IMR-90 clones. Number of cell divisions was calculated from total expansion data, using simulated culture viabilities ranging from 85-100%, and adjusting for uncounted high-VAF variants arising early during clonal expansion (see Materials and Methods). An average viability of 98.1%, equating to an estimated growth rate of 1.974 population doublings per cell division, was observed when the clones were harvested. (F-G) sSNV and sINDELs per bp detected using different combinations of WGS and NS libraries. Bars labeled (*a*), (*b*), and (*c*) correspond to the variant types outlined in Fig. 5A. Ideally, the sum of variants in *a* and *b* should equal *c*. n = 1 WGS library for bulk IMR-90s

We then estimated the mutation rate per cell division of each clone from *in vitro* expansion data, using simulated viabilities ranging from 90-100%. Because the NS pipeline screens out mutations with VAFs > 1% within the matched normal, we assumed that mutations stemming from the first 6.64 clonal divisions were not reported, and corrected the estimated number of cell divisions accordingly. Under these assumptions, we found that *de novo* SNV rates per cell division varied from 30.9 to 38.7 per diploid genome (**Fig. 5D**). Considering our observed culture viability of 98.1%, equating to a growth rate of 1.97 doublings/generation, we estimated a sSNV rate per cell division of 37.6 per diploid genome. This result is substantially larger than our previous calculation of 16.7 sSNVs/cell division, derived from estimates of post-zygotic doublings^23^. While our more robust cell division estimate may partially account for this discrepancy, these data suggest that human fibroblasts exhibit a higher somatic mutation rate *in vitro* than *in vivo*. We also found adjusted sINDEL mutation rates of 6.2 to 7.8 per diploid genome per cell division (**Fig. 5E**), yielding an estimate of 7.6 sINDELs/cell division at a viability of 98.1%.

Lastly, we reasoned that post-zygotic mutations found within a clone (*c*) could arise from one of two sources: somatic variants inherited from the original parent cell (*a*), or *de novo* mutations that occurred during clonal expansion (*b*) (**Fig. 5A**). After quantifying (*b*) during our initial analysis (**Figs. 5B-C**), we calculated (*a*) by comparing clone WGS libraries to bulk WGS data from the original IMR-90 population (**Fig. S4A-B**). We also used bulk WGS data as a matched normal for clone NS libraries to directly estimate (*c*) (**Fig. 5A**, *bottom right*). As a result, by assessing whether the sum of (*a*) and (*b*) was consistent with (*c*), we could validate the fidelity of our mutation calling approach. We found that for sSNVs, after correcting (*b*) for clonal high-VAF mutations, the sum of (*a*) and (*b*) averaged 15% less than (*c*) across the three clones (**Fig. 5F**). Despite this minor discrepancy, potentially resulting from selection effects or differences in variant calling criteria, we concluded that the NS estimates of sSNV burden were largely consistent with each other, as well as with clone WGS data. Although estimates of (*a*) and (*b*) for sINDELs differed from (*c*) by an average of 49% (**Fig. 5G**), the cumulative burden of (*a*) and (*b*) across all three clones differed from our cumulative estimate of (*c*) by only 3.6%. Thus, while individual measurements of sINDEL burden may be disrupted by technical artifacts, NS can detect sINDELs with relative consistency across multiple technical replicates. For both sSNVs and sINDELs, variants in (*b*) consistently accounted for more than half of the mutation burden in (*c*), despite accumulating over only 30 days. Our estimates of (*b*) and (*c*) were both derived from NS data, suggesting that our earlier observations, including reduced sSNV-to-sINDEL ratios and elevated sSNV rates per cell division, resulted from *in vitro* clonal expansion rather than sequencing discrepancies between NS and clone WGS. Overall, we concluded that somatic mutation burdens detected by *in silico*-adjusted NS are robust, informing reliable estimates of mutation rate per cell division in human fibroblasts.

### Mutational signatures and hotspots differ by method

While total mutation burden is sufficient to estimate the somatic mutation rate, the locus-specific context of somatic mutations can provide critical information about mutagenesis and selection events experienced by a cell population. To this end, we analyzed mutational signatures and hotspots from lymphoblastoid benchmark lines and IMR-90 fibroblasts. We first calculated the cosine similarity between each sample’s sSNV trinucleotide profile and the COSMIC Single Base Substitution (SBS) signatures^24,25^ (**Fig. S5**). We observed broad enrichment of SBS5, a clock-like signature associated with age^26^; SBS18, which has been linked to reactive oxygen species; and SBS40a-c, which have an unknown etiology.

Next, to assess whether somatic mutation patterns differed by cell type or pipeline, we isolated discrete mutational signatures from our sample pool using SigProfilerExtractor^27^. We identified two dominant mutational signatures within our samples: SBS96A, which included a high volume of C>A transversions (**Fig. 6A**, top), and SBS96B, which exhibited more C>T transitions (**Fig. 6A**, bottom). We expected these signatures to reflect cell type-specific differences between lymphoblastoid cell lines and IMR-90 fibroblasts. Surprisingly, however, after comparing the relative contribution of SBS96A and SBS96B to each sample’s mutation profile, we found that signature distribution differed by sequencing method in lymphoblastoid cell lines (**Fig. 6B**). While NS profiles of G1 and G2 aligned closely with SBS96A, mutations in SCMDA and WGNS samples were overwhelmingly attributed to SBS96B. Conversely, both NS and clone WGS data supported SBS96A as the dominant mutational signature in IMR-90 fibroblasts. Notably, neither mutational signature appears to be an artifact: SBS96B does not match mutational signatures observed in other SCMDA studies^28,29^, while SBS96A is supported by data from single-cell clones. As a result, it is likely that lymphoblastoid cell lines harbor an elevated number of C>T transitions, which are detectable by SCMDA and WGNS, but not by NS, possibly due to the specific restriction enzyme used by the latter for genome fragmentation.

**Figure 6.**
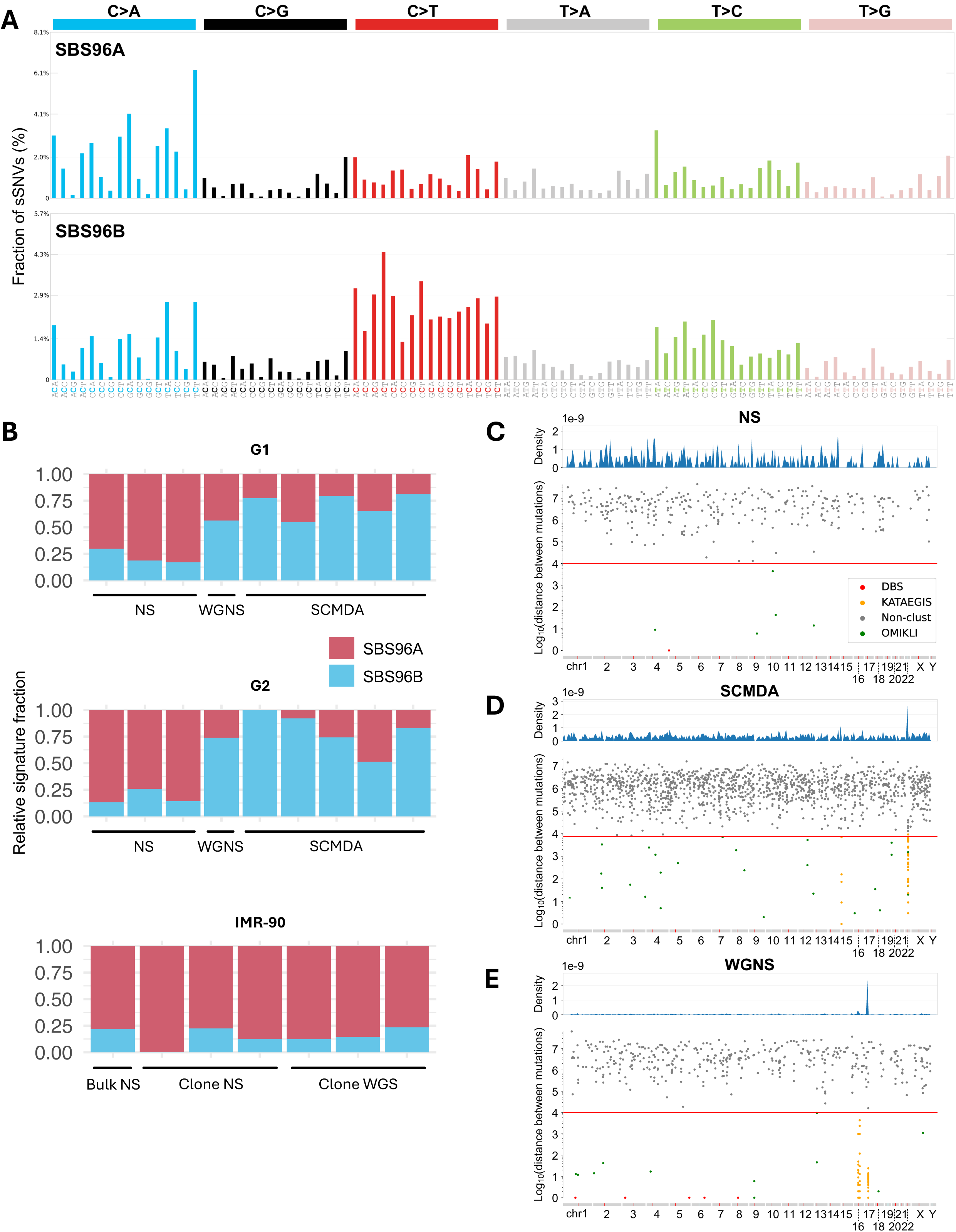
Mutational signatures and hotspots detected by single-cell and single-molecule sequencing methods. (A) sSNV trinucleotide signatures detected in pooled G1, G2, and IMR-90 samples. Signatures were extracted by non-negative matrix factorization using SigProfilerExtractor. n = 5 G1 SCMDA, 3 G1 NS, 1 G1 WGNS, 5 G2 SCMDA, 3 G2 NS, 1 G2 WGNS, 3 IMR-90 clone WGS, 1 IMR-90 bulk WGS, and 3 IMR-90 clone NS samples. (B) Relative contribution of SBS96A and SBS96B to the individual mutation signatures of each sample. (C-E) Representative genome-wide distribution of sSNVs in NS, SCMDA, and WGNS samples. Hotspots, denoted as kataegis sites, were identified by SigProfilerClusters. n = 1 sample per pipeline. Sample IDs: G2_16 (NS), G12878_5 (SCMDA), and G1_NS2_25 (WGNS).

Lastly, we used SigProfilerClusters^30^ to identify mutation hotspots within each sample, defined as dense clusters of sSNVs, e.g., “kataegis”^31^. Consistent with mutational signature data, we found that the genome-wide distribution of mutation hotspots differed by sequencing method. Although none of the pipelines identified hotspots in IMR-90 fibroblasts (**Figs. 6C & S6**), SCMDA and WGNS detected statistically significant kataegis in both lymphoblastoid lines (**Figs. 6D-E, S7 & S8**). Unlike in previous studies of B cell mutational spectra^28^, the ten hotspots identified by WGNS were concentrated near chromosome centromeres (**Fig. S9A**), regions characterized by poor sequencing coverage (**Fig. 2E**). As a result, we attributed these sSNV calls to technical artifacts introduced by the WGNS pipeline, which are preventable by raising the confidence threshold for primary read alignment (AS-XS) to the same level used for NS. Conversely, while only detected in 3 out of 10 cells, hotspots identified by SCMDA were enriched at immunoglobulin genes (**Fig. S9B**), reflecting somatic hypermutation (SHM) during B cell development^32^. Notably, we found that C>T transitions far outnumbered other sSNVs in SCMDA hotspot regions (**Table S5**), suggesting that sensitivity toward SHM could explain differences in mutational signature composition between NS and SCMDA. Although we did not observe preferential mutagenesis at TT dinucleotides, which we had previously linked to SHM *in vivo*^28^, it is possible that *in vitro* SHM following EBV infection exhibits a preference for alternative DNA repair pathways^33–35^, resulting in distinct mutational spectra.

Overall, we found that the choice of sequencing method can substantially influence observed mutational signatures and hotspots, with SCMDA detecting evidence of lymphoblastoid SHM that was absent in NS and WGNS samples. These results strongly reinforce the need for standardized benchmarks and error correction techniques to facilitate robust somatic mutation characterization.

## Discussion

In this study, we developed both *in vitro* and *in silico* approaches to benchmark somatic mutation calling in standard cell lines using state-of-the-art single-cell and single-molecule DNA sequencing methods. We demonstrated that SCMDA, NS, and WGNS can detect sSNVs and sINDELs with high accuracy at the single-cell level. However, each method has limitations: SCMDA is less cost-efficient; NS exhibits bias toward certain genomic regions and sequence contexts; and sINDEL calling with NS and WGNS remains hindered by a relatively high rate of artifacts. Our approaches and results using standard cell lines provide an essential benchmark for ensuring the accuracy of somatic mutation calling in noncancerous cells. These technologies advance our fundamental understanding somatic mosaicism in normal tissues, which may contribute to many chronic diseases beyond cancer, as well as to the functional decline associated with aging^2^.

Beyond overall accuracy, our analyses revealed that the choice of sequencing method can substantially influence the observed mutational landscape. For example, SCMDA detected clusters of C>T transitions in immunoglobulin loci, consistent with somatic hypermutation in B cells, whereas NS failed to capture these hotspots. While this may simply have resulted from a low incidence of somatic hypermutation within the G1 and G2 populations, reducing the likelihood of detecting these events in a bulk DNA library, mutational signatures in NS samples also differed from those detected by SCMDA and WGNS. This raises the possibility of restriction enzyme-driven coverage bias, which, by preventing detection of somatic variants near cut sites, may have affected the observed mutational signatures. WGNS, while providing broad genomic coverage, produced hotspots near centromeres that were likely composed of technical artifacts, which could be eliminated by optimizing computational pipelines. Such method-specific differences in mutational signatures and hotspot distribution underscore the risk of misinterpreting biological patterns, particularly in studies of aging, environmental exposures, or disease associations, if sequencing biases are not properly accounted for.

To demonstrate the broader utility of our benchmark, we used an *in silico*-adjusted pipeline to estimate the somatic mutation rate per cell division in early-passage IMR-90 human fibroblasts. We observed an average accumulation of ∼37.6 sSNVs and ∼7.6 sINDELs per diploid genome per division *in vitro*, substantially higher than previous estimates of mutation rate in primary fibroblasts based on post-zygotic divisions^23^. This elevated rate may have resulted from primary cells in culture experiencing increased replication stress, oxidative damage, and potentially altered selection pressure^36^. These findings emphasize that mutation rates measured *in vitro* may not directly reflect *in vivo* physiology, and that culture-specific mutational processes should be considered when using primary cell models to study somatic mosaicism, aging, or disease.

Although our benchmark provides a quantitative means to compare single-cell and single-molecule pipelines, the pTP and pFP estimates remain imperfect. In both G1 and G2 cell lines, the adjusted mutation burdens per base pair still differed between single-cell (SCMDA) and single-molecule (NS/WGNS) data, suggesting that our adjustment does not fully reconcile platform-specific biases. One plausible explanation is that we overestimated the pFP component: in practice, a substantial fraction of calls classified as pFPs are likely to be genuine somatic mutations present in the assayed sample but absent from the mismatched “normal” genome. This “misclassification” by design would artificially inflate the estimated false-positive rate and consequently lead to underestimation of the true mutation burden, disproportionately affecting the adjusted estimates across the three methods.

Overall, our study highlights the substantial impact of technology-specific biases on somatic mutation detection and interpretation, and establishes a foundation for standardized, cross-platform-comparable analyses of somatic mosaicism in normal human tissues.

## Data availability

Raw sequencing data will been deposited in the NCBI Sequence Read Archive (SRA) before publication.

## Acknowledgement

This work was supported by the American Federation for Aging Research (the Sagol Network GerOmic Award for Junior Faculty to L.Z.), the U.S. National Institutes of Health (P01 AI172501 to X.D., U19 AG056278 to X.D., and R35 GM159832 to L.Z.).

## Materials and Methods

### Sample Preparation

The NIST standard lymphoblastoid lines GM12878 (G1) and GM24385 (G2), as well as their purified genomic DNA (NA12878 and NA24385, respectively), were acquired from the NIGMS Human Genetic Cell Repository at the Coriell Institute for Medical Research. Both GM12878 and GM24385 were cultured in complete RPMI 1640 containing 15% FBS, 100 IU/mL penicillin, and 100 μg/mL streptomycin. Early-passage IMR-90 human fibroblasts were obtained from the NIA Aging Cell Culture Repository at the Coriell Institute. IMR-90 cells were grown in EMEM containing 10% FBS, 100 U/mL penicillin, and 100 μg/mL streptomycin. All cultured cells were maintained at 37 °C with 5% CO_2_ and 3% O_2_.

### Single Cell Collection and Clonal Amplification

For single-cell whole-genome sequencing, single cells from GM12878 and GM24385 were collected with the CellRaft system (Cell Microsystems, Research Triangle Park, North Carolina). Briefly, a CellRaft array was rinsed with PBS and coated with 2% gelatin in water (G1393-100ML; Sigma-Aldrich) at 37 °C for 1 hour. After removing excess gelatin solution, approximately 7,000 cells in 4 mL of complete RPMI 1640 medium were added to the array and incubated at 37 °C for 3 hours under 3% O₂ and 5% CO₂. Floating cells were removed by washing twice with 1 mL of fresh complete medium. Finally, 3 mL of fresh complete medium was added to the array. Single rafts containing one cell were individually transferred into 0.2 mL PCR tubes containing 2.5 μL PBS, then immediately stored at −80 °C.

For fibroblast clones, IMR-90 clones were also generated by seeding a single cell suspension on a CellRaft array. Because IMR-90 fibroblasts are adherent cells, gelatin coating was not required for their array. All other steps for preparing the samples were the same as those used for lymphoblastoid cells. Single rafts that possessed one cell after 24 hours were monitored for another 5 days, at which point colonies of 8-32 fibroblasts were picked. After an additional 25 days, the expanded clones, numbering 5.7-7.8×10^6^ cells, were collected and frozen at −80 °C in EMEM containing 10% FBS and 10% DMSO.

### Bulk Genomic DNA Extraction

DNA from cultured lymphoblastoid lines, bulk IMR-90s, and IMR-90 clones were isolated using either a GeneJET Genomic DNA Purification Kit (Thermo Fisher Scientific, Waltham, Massachusetts) or a Quick DNA/RNA Microprep Plus Kit (Zymo, Irvine, California). Purified genomic DNA samples, including those obtained from Coriell, were assessed for shearing and contamination using gel electrophoresis.

### Library Preparation for Single-Cell and Bulk Sequencing Pipelines

Single-cell multiple displacement amplification (SCMDA) was performed on frozen GM12878 and GM24385 single cells as previously described^17^. Genomic DNA samples from both cultured lymphoblastoid cell lines, bulk IMR-90s, and IMR-90 clones were prepared for standard whole-genome sequencing (WGS) using a NEBNext Ultra II FS DNA Library Prep Kit for Illumina (New England Biolabs, Ipswich, Massachusetts). Samples of purchased lymphoblastoid genomic DNA, as well as DNA from all IMR-90 populations, were prepared using a Nanorate Sequencing (NanoSeq or NS) protocol adapted from Abascal et al.^12^, with NEBNext Ultra II Ligation Module (Cat. No. E7595) replacing the original ligation mix to improve ligation efficiency. Samples of purchased lymphoblastoid genomic DNA were also subjected to whole-genome NanoSeq (WGNS) library preparation as described^18^.

### Library QC and Sequencing

Libraries were subjected to QC using Qubit (Invitrogen) and Bioanalyzer (Agilent). Samples were then sequenced with the Illumina NovaSeq S4 system or NovaSeq X Plus System at Novogene using 150-bp paired-end reads.

### Read Alignment and Variant Calling Methods

NS, and WGNS: As described^12^, raw NS data were first pre-processed, including trimming adapter sequences and extracting the duplex barcodes from the fastq files, added them to the fastq header of each read. Then reads were mapped to the human reference genome (version: GRCh38) using BWA MEM^37^. After alignment, A read bundle tag was appended to each read pair of a BAM to determine which reads were PCR duplicates, by using bamsormadup from biobambam2^38^. With bamaddreadbundles^12^, optical duplicates and unpaired mates were filtered. All the samples passed the efficiency check^12^.

The NS analysis requires a matched normal to distinguish somatic mutations from germline SNPs, thus we performed deep WGS of GM12878 and GM24385 as the matched normal. Using the wrapper script^12^, a coverage histogram was computed for the alignment file (bam file). SNP and noise sites were marked in the output VCF file. For NS and WGNS, we applied different parameter setting suggested by the original papers in the variant calling procedure^12,18^.

Specifically, the computational pipelines for NS and WGNS include a series of parameters that establish thresholds for somatic variant calling, including a minimum primary alignment score minus secondary alignment score (AS-XS; *a*) and a maximum VAF within the matched normal (*v*). In all cases where we performed NS or WGNS, we used the default parameters for both pipelines (NS: *a* = 50, *v* = 0.01; WGNS: *a* = 10, *v* = 0.1). Abascal et al.^12^ and Lawson et al.^18^ also provide two genome masks, containing common SNPs and recurrent technical artifacts, to reduce the false-positive rate of NS and WGNS. Because these masks selectively eliminate germline variants, we did not apply either mask when calculating pFP and pTP rates (**Fig. 3A-E**), including for our *in silico* adjustments (**Fig. S3**; **Table S3**). Since pTP rates have a much greater impact on our *in silico* adjustments than pFP rates (compare **Figs. 3D-E** to **Fig. S3**), the inclusion or exclusion of masks, which greatly reduce pFP calls (**Figs. 3B-C** vs **Table S3**) while minimally impacting coverage, should otherwise have little effect on the magnitude of our adjustments. For our *in vitro* experiments using 1:000 mixtures (**Figs. 3G-H**), we excluded the SNP mask when calculating mutation burden, since it removed nearly all mixed germline SNPs (data not shown). We also excluded the noise mask when calculating mixed INDEL burden, since the noise mask disproportionally eliminated germline INDEL sites (data not shown). In all other experiments (**Figs. 4&5**), both masks were used to calculate raw mutation burden (**Table S3**), which we then multiplied by our adjustment factor to yield an adjusted mutation burden.

scWGS, clone WGS, and bulk WGS: Raw sequence reads were subjected to quality control using FastQC and aligned to the human reference genome (GRCh38) using BWA MEM^37^. PCR duplicates were marked and removed using GATK^39^. Base-pair score quality of aligned reads was recalibrated using GATK. SNVs and INDELs observed in a cell or clone but not present in the corresponding bulk DNA were called by comparing the aligned sequences of the cell or clone to the bulk using SCcaller (version 2.0)^17^ with default options.

### Mutation burden per base pair and rate per cell division

For NS or WGNS,

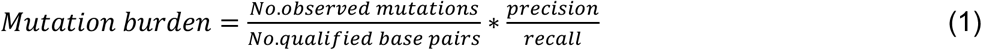

For scWGS, because the cells are of diploid genomes,

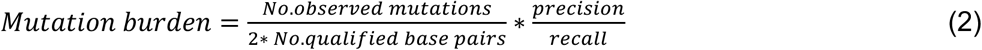

In the above,

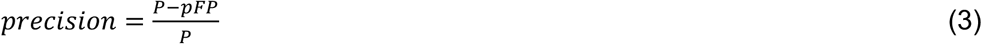

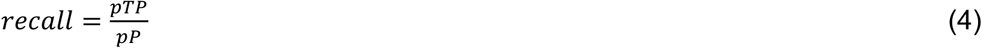

where pTP, pFP, pP and P are pseudo-True Positive, pseudo-False Positive, pseudo-Positive, and Positive calls respectively, obtained from heterozygous germline SNPs or INDELs by comparing NS, WGNS, or scWGS data of sample A (e.g., G1) to bulk WGS data of sample B (e.g., G2).

To estimate mutation rates per cell division, the adjusted per-base burden was scaled to a genome-wide value by multiplying by the haploid genome size. The resulting number of mutations was divided by the mitotic history of the clone, which was estimated under an exponential growth model starting from a single progenitor cell. The number of divisions were inferred from culture duration and cell doubling times. Dividing the corrected genome-wide mutation count by the estimated number of mitoses yielded the average number of new mutations per cell division.

---

**Figure S1.**
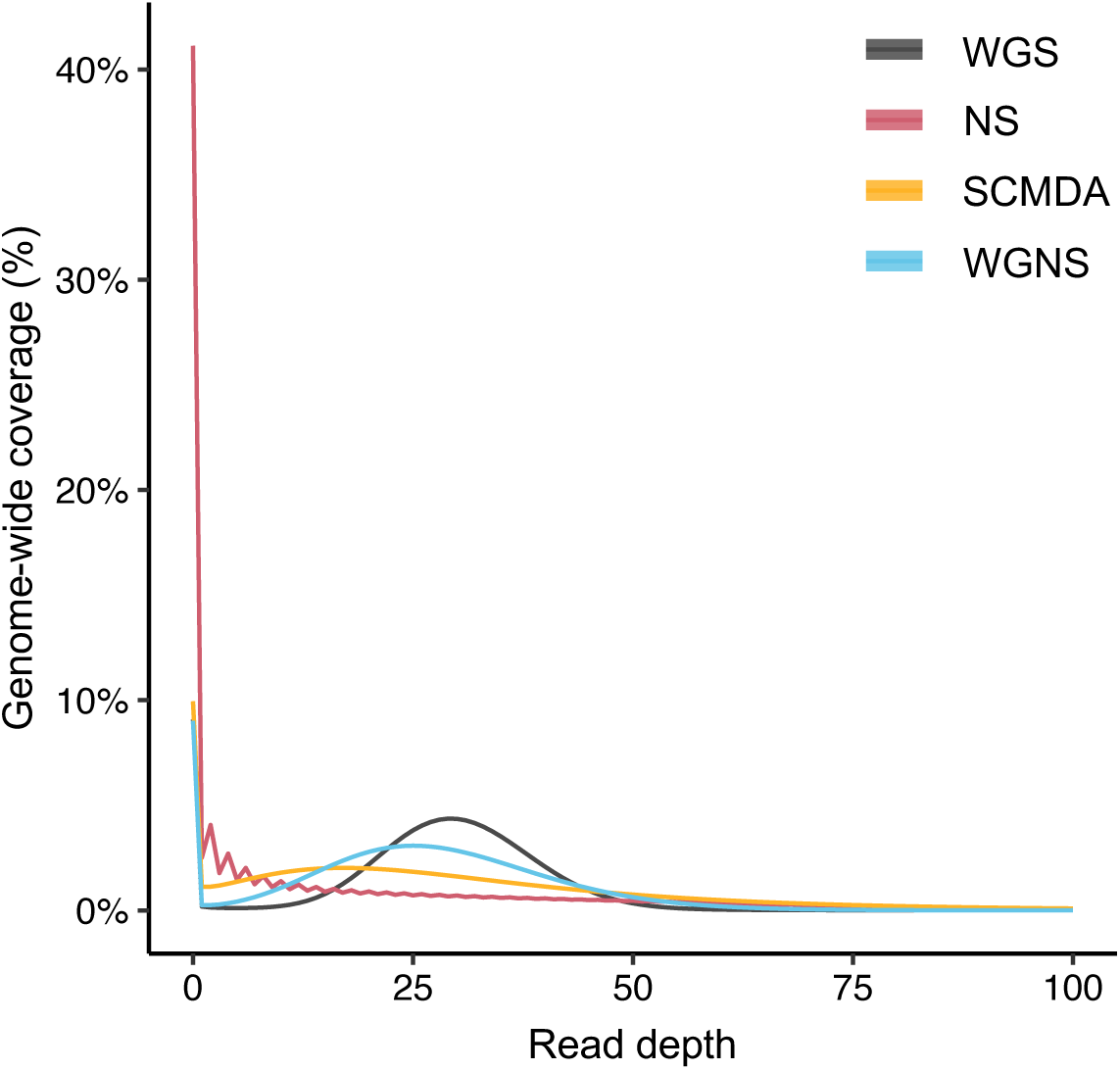
Coverage-depth distribution for WGS, NS, SCMDA, and WGNS. Data are equivalent to that displayed in Fig. 2C, but with the inclusion of coverage fractions at a read depth of zero. n = 6 WGS, 10 NS, 10 SCMDA, and 2 WGNS samples.

**Figure S2.**
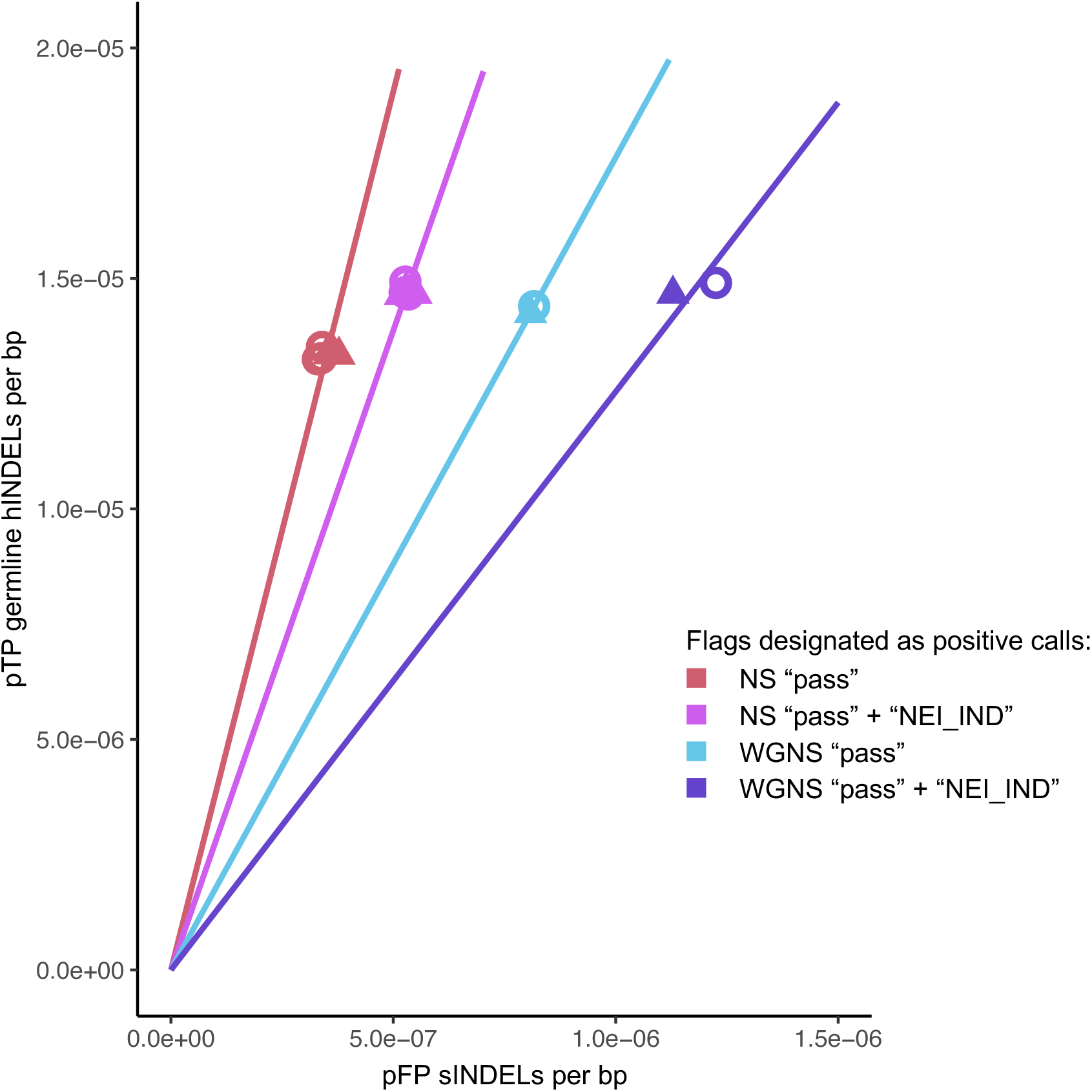
pFP sINDEL and pTP hINDEL rates per bp in NS and WGNS samples following inclusion of “NEI_IND”-marked variants. Samples and methodology are identical to those enumerated in Fig. 3E. n = 3 NS, 5 SCMDA, and 1 WGNS samples per cell line. During variant calling, the default NS and WGNS pipelines identify INDEL-rich sites within the matched normal, flagging putative somatic variants at these loci as “NEI_IND”. To validate the utility of our *in silico* approach as a tool for pipeline optimization, we assessed whether exclusion of NEI_IND calls enhances variant calling accuracy. Despite a slightly higher pTP rate, alternate NS and WGNS pipelines that included these variants exhibited a disproportionally high pTP rate, supporting the decision by the original authors to eliminate INDEL-rich sites. Thus, our *in silico* test can rapidly assess the impact of parameter settings on pipeline performance, facilitating pipeline development.

**Figure S3.**
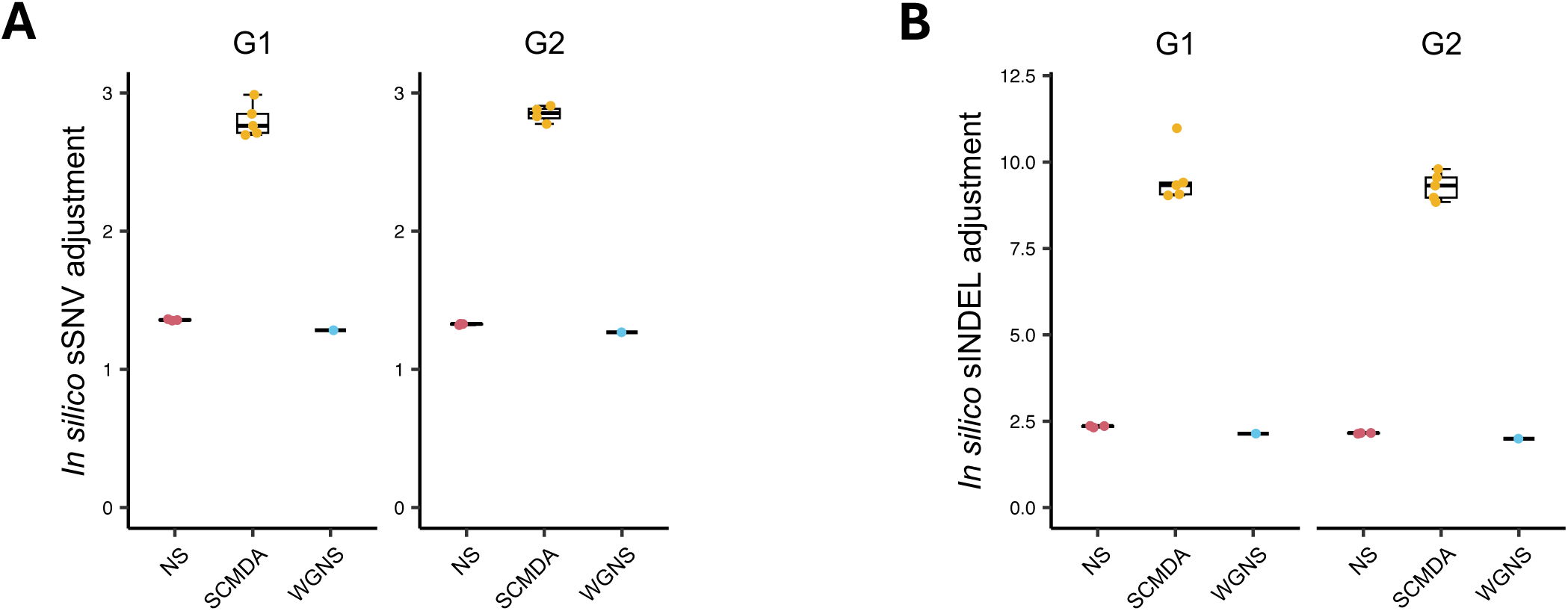
(A-B) sSNV and sINDEL adjustment factors for NS, SCMDA, and WGNS samples. Adjustment factors were calculated based on pFP and pTP rates derived from *in silico* testing (see Materials and Methods). Each point represents a single sample, while box plots display the mean and quartile statistics for each sample group. n = 3 NS, 5 SCMDA, and 1 WGNS samples per cell line for G1 and G2.

**Figure S4.**
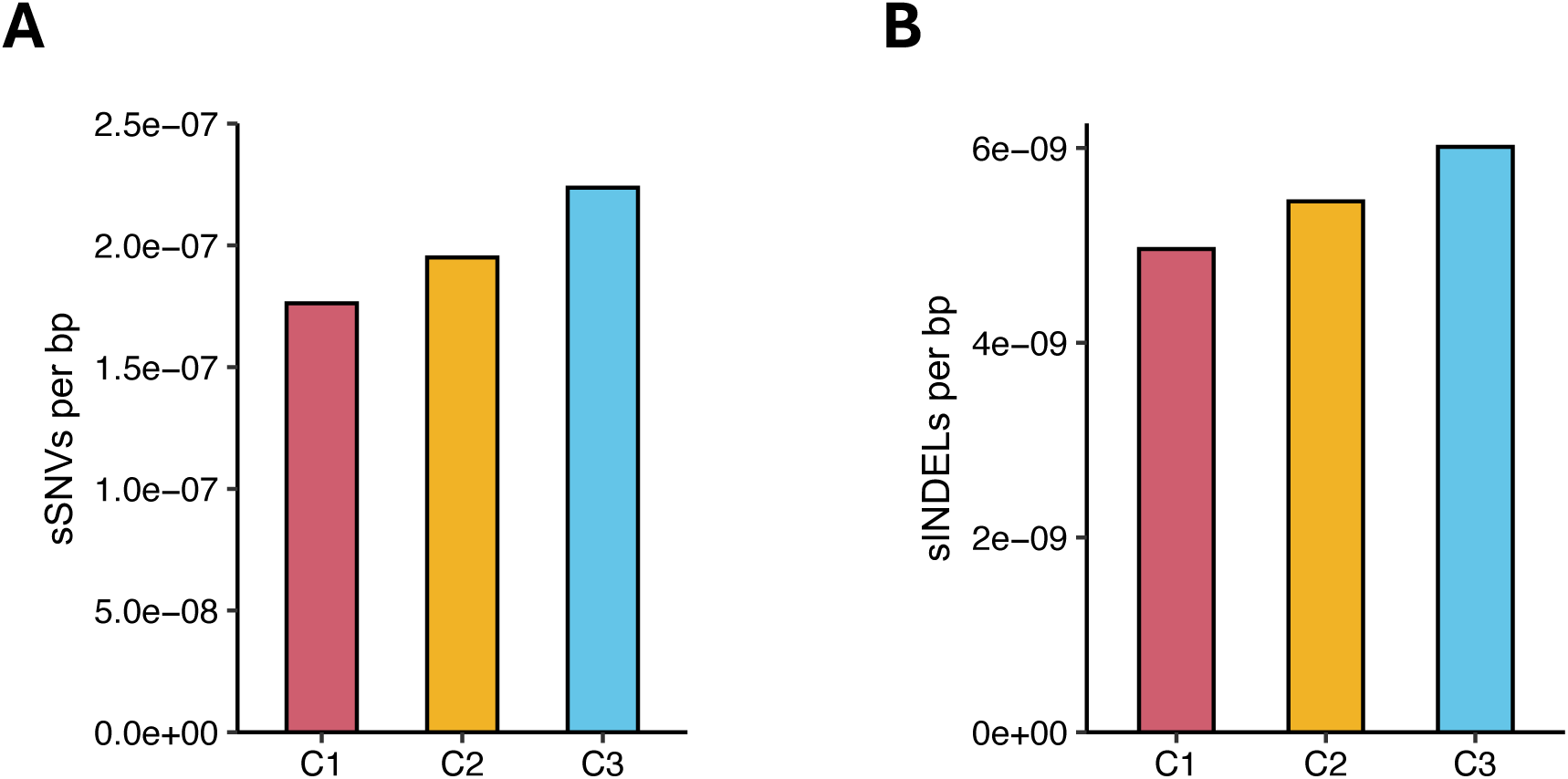
(A-B) Parent cell sSNV and sINDEL mutation burdens detected in IMR-90 clones. Mutation burdens were calculated by screening clone WGS libraries against bulk WGS data (Fig. 5A, *top right*). A single WGS library was constructed for each clone, while one additional library was constructed for bulk IMR-90s. n = 3 clones. Note: the cells subjected to bulk and clone WGS were descended from the same initial population, but did not possess the same population doubling level.

**Figure S5.**
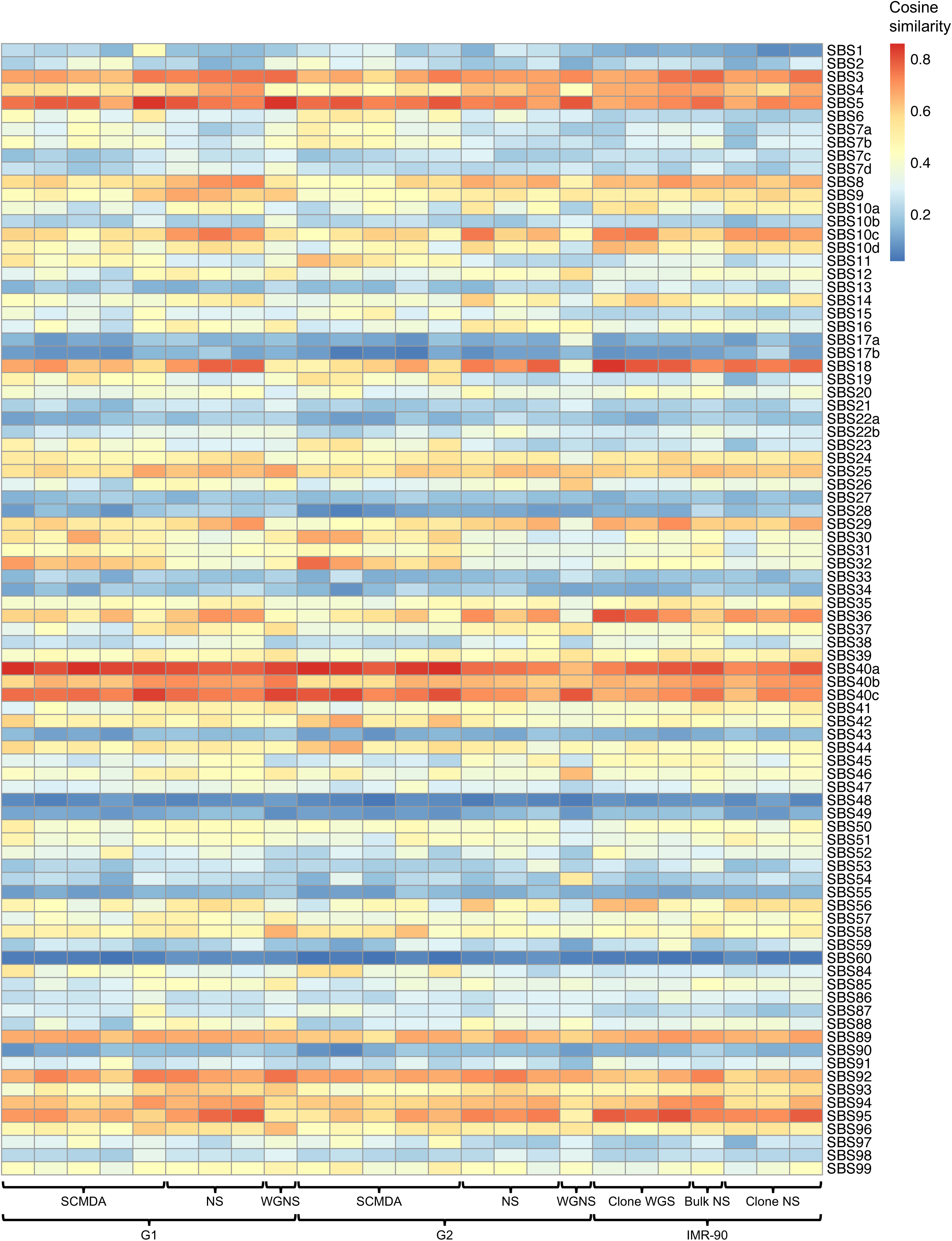
Cosine similarity matrix for sSNV trinucleotide signatures. Cosine similarities were calculated between observed mutational spectra and COSMIC v3.4 reference signatures. A heatmap of similarity scores was generated, without clustering, to highlight signature correspondence by cell type and method. n = 5 G1 SCMDA, 3 G1 NS, 1 G1 WGNS, 5 G2 SCMDA, 3 G2 NS, 1 G2 WGNS, 3 IMR-90 clone WGS, 1 IMR-90 bulk WGS, and 3 IMR-90 clone NS samples.

**Figure S6.**
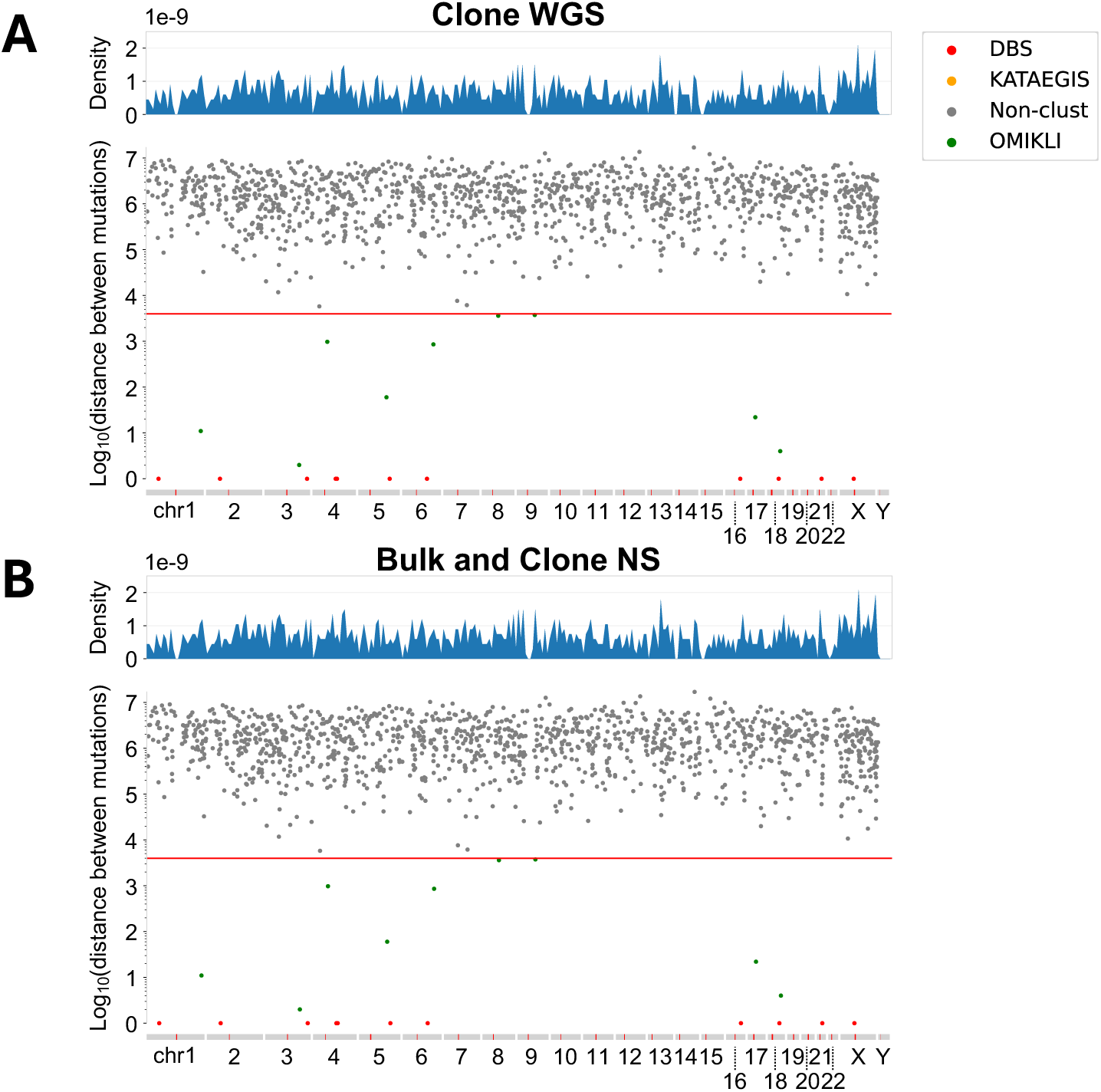
(A) Genome-wide distribution of sSNVs in IMR-90 clone WGS samples. Variants were pooled from three distinct IMR-90 clones. Clustering was performed using SigProfilerClusters. (B) Genome-wide distribution of sSNVs in bulk NS and clone NS samples. Variants were pooled from one bulk NS and three clone NS samples.

**Figure S7.**
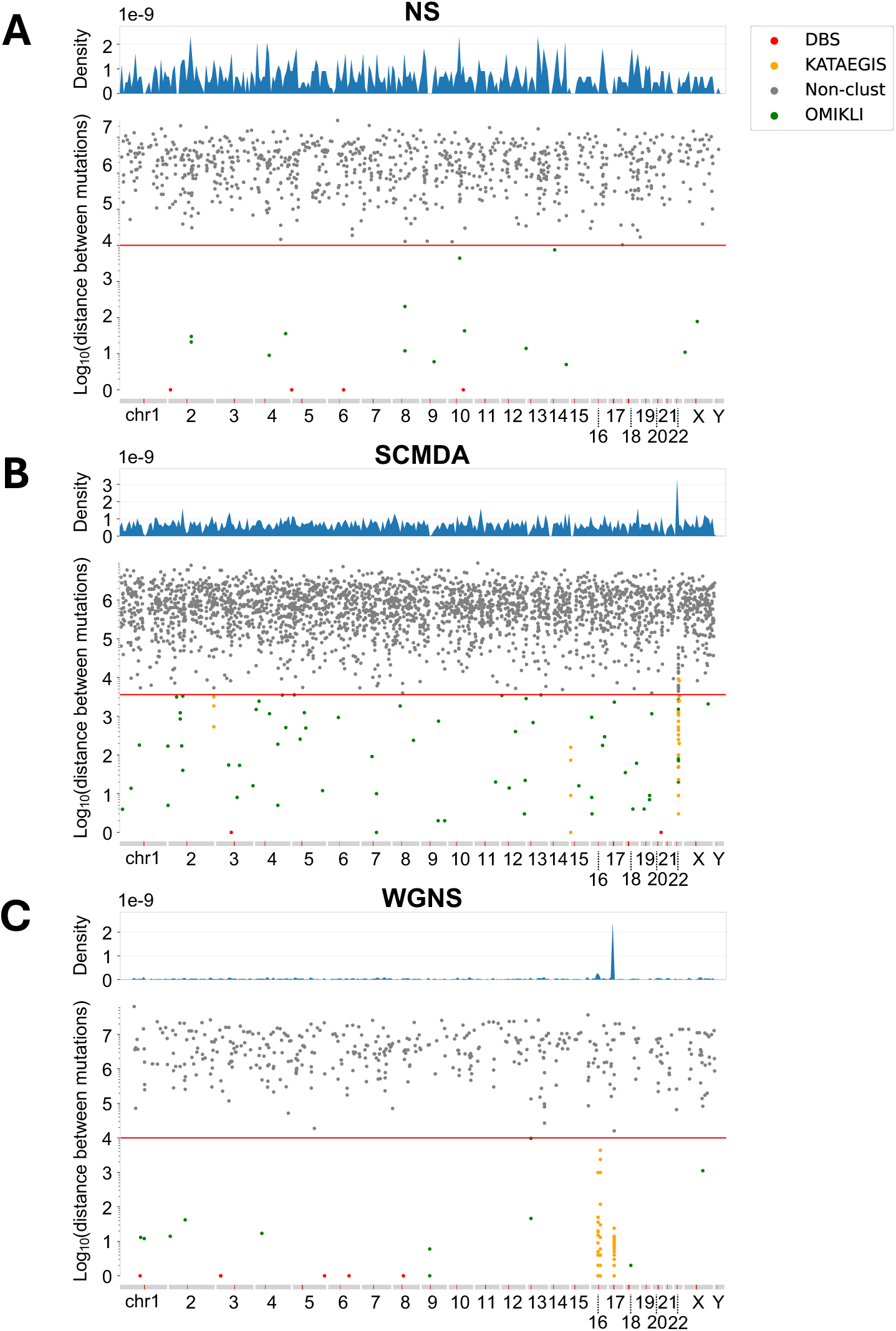
(A-C) Genome-wide distribution of sSNVs in G1 NS, SCMDA, and WGNS samples. All replicates within each pipeline were pooled. Clustering was performed using SigProfilerClusters. n = 3 NS, 5 SCMDA, and 1 WGNS samples.

**Figure S8.**
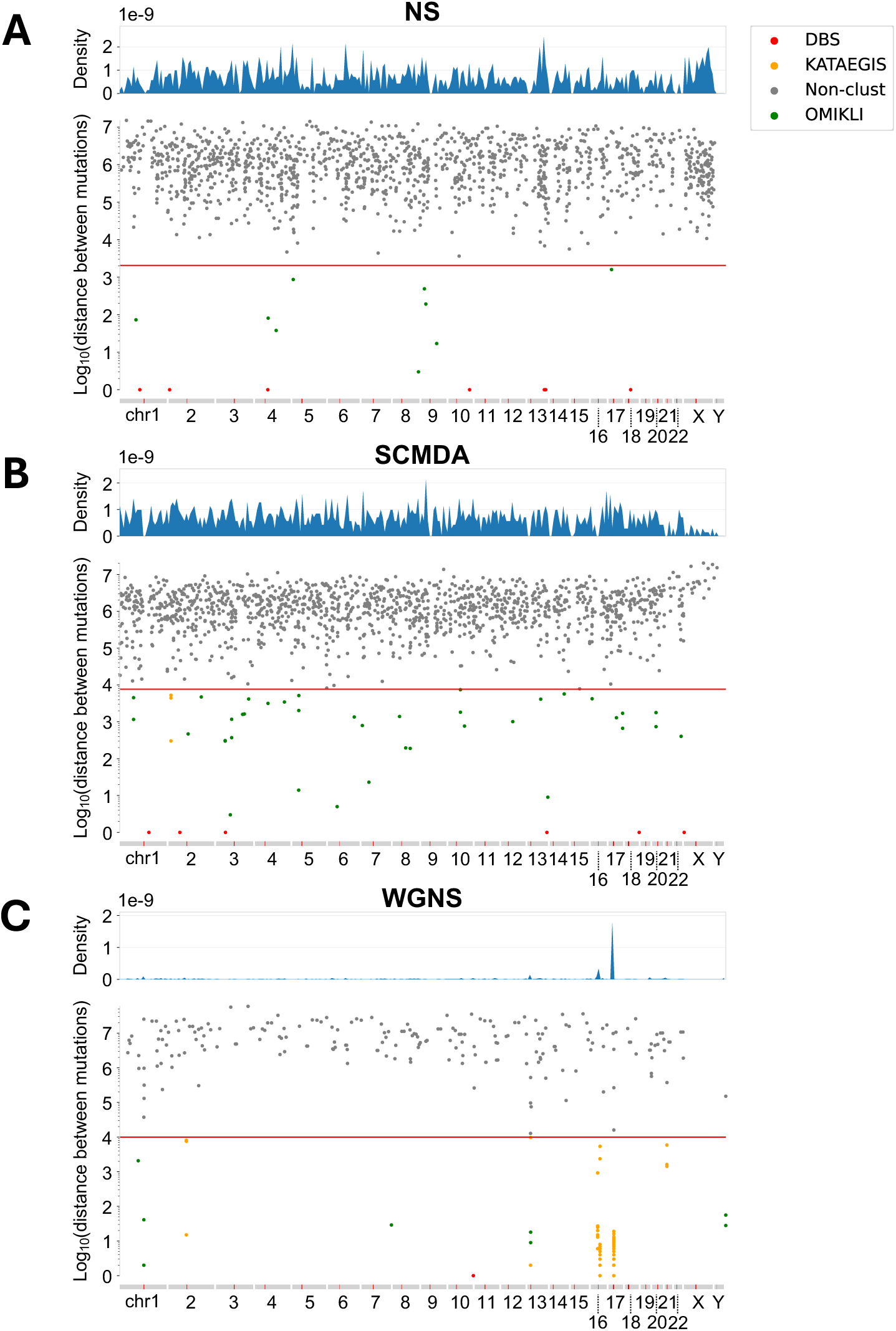
(A-C) Genome-wide distribution of sSNVs in G2 NS, SCMDA, and WGNS samples. All replicates within each pipeline were pooled. Clustering was performed using SigProfilerClusters. n = 3 NS, 5 SCMDA, and 1 WGNS samples.

**Figure S9.**
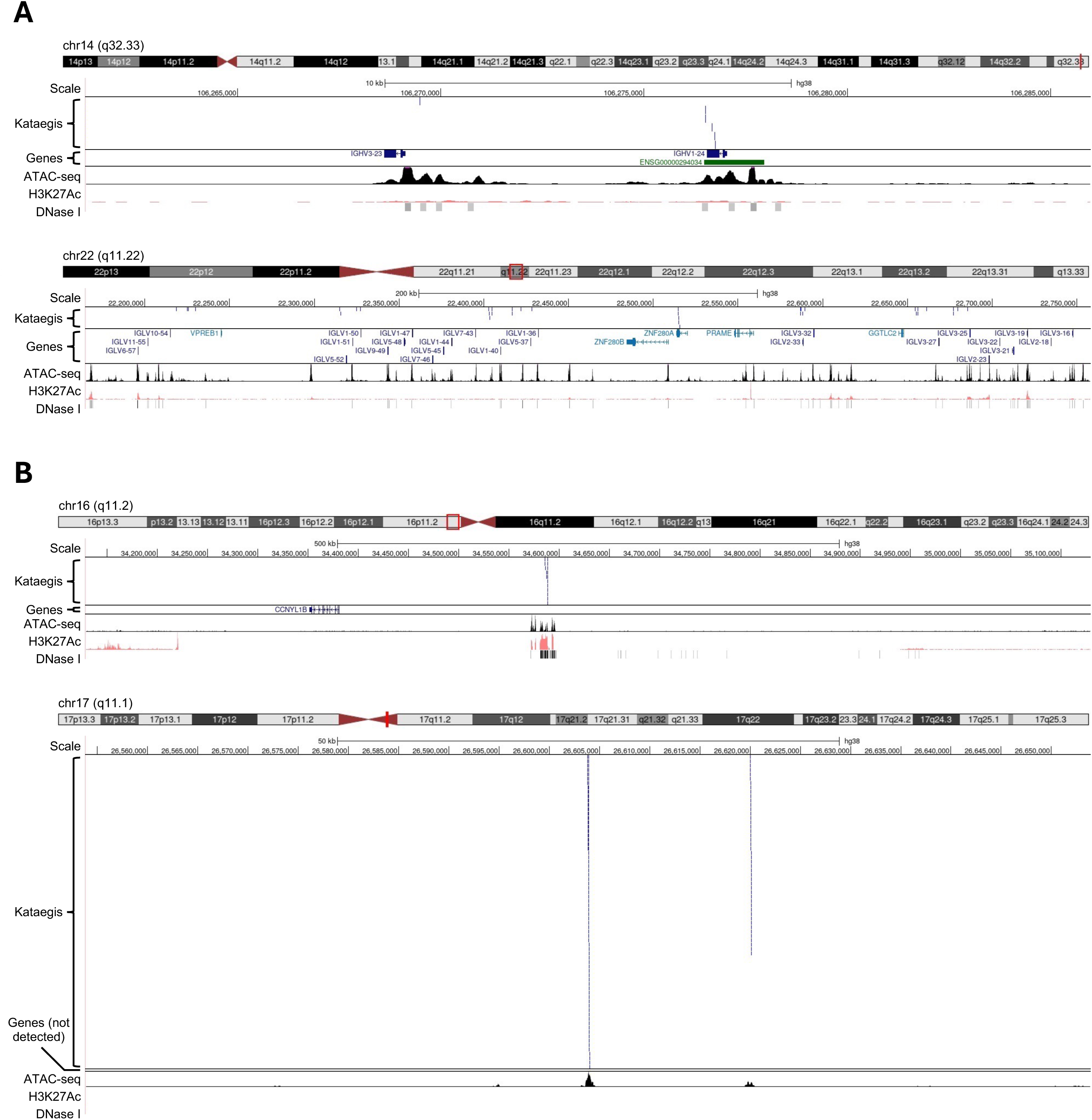
Characteristics of sSNV hotspots in SCMDA and WGNS samples. (A) Representative view of kataegis sites in chromosomes 14 (*top*) and 22 (*bottom*) of SCMDA samples. Loci were visualized using the UCSC genome browser, including gene annotations, ATAC-seq data, H3K27Ac sites, and DNase I hypersensitivity sites. (B) Representative view of kataegis sites in chromosomes 16 (*top*) and 17 (*bottom*) of WGNS samples.

